# Intestelligence: A pharmacological neural network using intestine data

**DOI:** 10.1101/2023.04.15.537044

**Authors:** Yusuke Watanabe, Hiroshi Ban, Nobuhiro Hagura, Yuji Ikegaya

**Affiliations:** Graduate School of Pharmaceutical Sciences, The University of Tokyo, Tokyo 113-0033, Japan; Institute for Advanced Co-creation studies, Osaka University, Osaka 565-0871, Japan; Center for Information and Neural Networks, National Institute of Information and Communications Technology, Suita City, Osaka, 565-0871, Japan; Institute for AI and Beyond, The University of Tokyo, Tokyo 113-0033, Japan

## Abstract

A neural network is a machine learning algorithm that can learn and make predictions by adjusting the strength of the connections between nodes. The sigmoid function is commonly used as an activation function in these nodes. This study explores the potential applicability of biological materials in the development of alternative activation functions. Inspired by the fact that acetylcholine induces intestinal contractions that follow a sigmoid function, we used pharmacological data obtained from guinea pig ilea in a layered neural network for image classification tasks. We found that the intestinal data-based neural network with the same structure as a conventional three-layer perceptron achieved an impressive classification accuracy of 85.7% ± 0.6% based on the MNIST handwritten digit dataset (chance = 10%). Additionally, the neural network was trained to determine whether objects in photographs collected from the internet were digestible, achieving an accuracy of 88.5% ± 0.9% (chance = 50%). Our approach highlights the potential applicability of intestine data in neural computations based on pharmacological mechanisms.

## Introduction

Neural networks (NNs), which were first introduced by Amari (1967) and later improved by Rumelhart et al. (1986), are mathematical models that are inspired by the structure and function of the central nervous system (CNS). A critical aspect of NNs is their nonlinearity, which is provided by activation functions that control the nodes’ outputs, similar to biological neurons.

Despite their important impact on neural network behavior, many activation functions have been proposed without theoretical justification (Glorot et al., 2011; Hahnloser et al., 2000; Jarrett et al., 2009; Nair & Hinton, 2010; Ramachandran et al., 2018; Misra, 2020). Therefore, the use of biologically verified activation functions, such as those derived from the enteric nervous system, may improve the performance of existing activation functions. The enteric nervous system shares many features with the central nervous system (Goyal & Hirano, 1996; Rao & Gershon, 2016) and may show promise for the development of unconventional neural network models.

The intestine, which is sometimes referred to as the “second brain” (Mayer, 2011; Obata & Pachnis, 2016; Rao & Gershon, 2016; Maniscalco & Rinaman, 2018), responds to external stimuli. Its smooth muscle undergoes reversible contractions in response to acetylcholine (ACh) in a dose-dependent manner (Burgen & Spero, 1997; Rao & Gershon, 2016). These contractions conform to a dose‒response curve that reflects the activation of ACh receptors and subsequent signal pathways; thus, contraction responses are macroscopic expressions of biochemical reactions. Therefore, intestinal contraction data may be a valuable resource for developing novel biologically inspired activation functions.

In this study, we developed an intestinal data-based neural network (INN) by incorporating pharmacological data based on intestinal contractions into a neural network model. We hypothesized that the INN model could perform classification tasks as effectively as neural networks based on the central nervous system.

## Materials and Methods

### Animal ethics

The intestinal tissues used in this study were surplus samples in a university student practical course that would otherwise have been discarded. Specifically, retired male breeder guinea pigs weighing 300–400 g (*n* = 14; SLC, Shizuoka, Japan) were used in our experiments. The animals were housed in a temperature-controlled room (22 ± 1°C) with a 12-hour light-dark cycle (light from 07:00 to 19:00) and given *ad libitum* access to food and water. All animal experiments were performed with the approval of the Animal Experiment Ethics Committee of the University of Tokyo (approval number: P24-1) and in accordance with the University of Tokyo guidelines for the care and use of laboratory animals. Our experimental protocols adhered to the Fundamental Guidelines for Proper Conduct of Animal Experiments and Related Activities in Academic Research Institutions (Ministry of Education, Culture, Sports, Science and Technology, Notice No. 71 of 2006), the Standards for Breeding and Housing of and Pain Alleviation for Experimental Animals (Ministry of the Environment, Notice No. 88 of 2006), and the Guidelines on the Method of Animal Disposal (Prime Minister’s Office, Notice No. 40 of 1995).

### Drugs

Acetylcholine chloride (C_7_H_16_ClNO_2_; Wako, Japan) was dissolved in double distilled water at concentrations of 2×10^−7^, 2×10^−6^, 2×10^−5^, 2×10^−4^, 2×10^−3^, 2×10^−2^, and 2×10^−2^ M.

### Intestinal contraction data acquisition

To isolate ileum tissue, guinea pigs were deeply anesthetized with 100 mg/kg pentobarbital (i.p.). The chest was opened, and the ileum was removed and washed with nutrient solution (in mM: NaCl 137, KCl 2.7, NaHCO_3_ 11.9, CaCl_2_ 1.8, MgCl_2_ 1.0, D-glucose 5.5) that was bubbled with 5% CO_2_/95% O_2_. The tissue was cut to a length of approximately 3 cm, and mesenteric arteries were removed. Two hooks were used to mount the tissue; one hook was inserted at one end of the tissue and fixed to the tip of an aeration tube, while the other was connected to an isotonic displacement transducer (45347, NEC San-ei Instrument, Tokyo, Japan). The transducer was connected to an amplifier (AS1202, NEC San-ei Instruments) and a flatbed recorder (FBR-251A, TOA Electronics, Tokyo, Japan), which was used to record the ileum tissue responses (Fig. 1A). Before starting the experiments, the ileum tissue strips were equilibrated, and a stable baseline recording was obtained at a resting tension of 0.5 g. Acetylcholine (ACh) solutions were then sequentially bath-applied to the organ bath at the following concentrations (in M): 1×10^−9^, 3×10^−9^, 1×10^−8^, 3×10^−8^, 1×10^−7^, 3×10^−7^, 1×10^−6^, 3×10^−6^, 1×10^−5^, 3×10^−5^, 1×10^−4^, 3×10^−4^, 1×10^−3^, and 3×10^−3^. Bath application was performed when a local maximum of the contraction reaction at the previous ACh concentration was confirmed. The ACh concentration [M] and intestine length [cm] were recorded, and these steps were repeated 2–3 times. The intestinal contraction rate [%] was calculated by averaging the contraction lengths over different trials and dividing the mean values by the maximum value. We collected the data for the dose‒response curves from 512 ileum samples.

**Figure 1.**
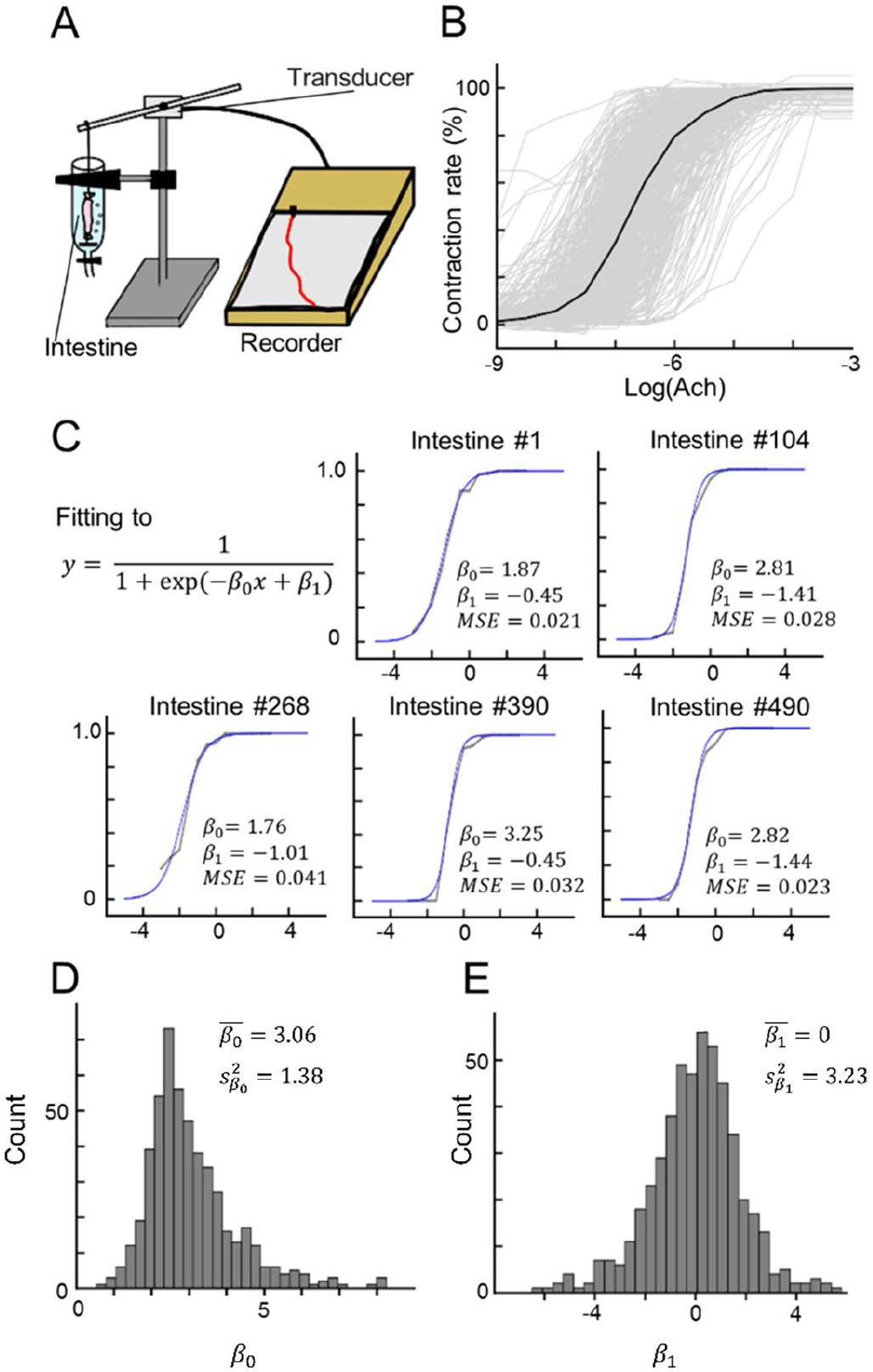
Dose‒response curves of ACh-induced contractions of the ileum. (A) Illustration of our experimental devices for measuring ileum contraction. (B) The dose‒response curves of 512 data points for contractions in response to bath application of ACh. Each *gray* line indicates a single data point, whereas the *black* line indicates the mean. (C) Five examples of sigmoid fitting of the dose‒response curves of ACh-induced ileum contraction. Note that β_0_ and β_l_ are fitting parameters. (D, E) Distributions of the fitting parameters β_0_ (D) and β_l_ (E).

### Sigmoid-curve fitting for the intestine contraction data at different acetylcholine concentrations

For each of the 512 data pairs, the ACh concentration was natural-log-transformed:

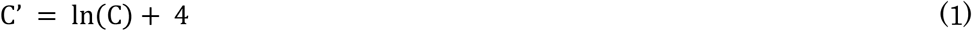

where C is the ACh concentration in the organ bath. The dose‒response curves were plotted as

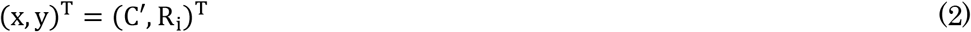

where R_i_ is the i^th^ intestine contraction rate [%] (i = 1, 2, …, 512; Fig. 1B). These curves were fitted by sigmoid functions:

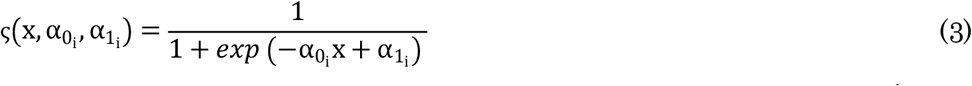

where 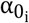 and 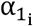 are parameters to fit the sigmoid function based on the i^th^ intestine data. The parameters 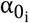 and 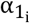 were optimized simultaneously using the MATLAB function nlinfit (MathWorks, Natick, MA, USA). The determined sigmoid functions were shifted by 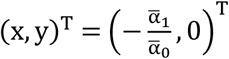 as follows:

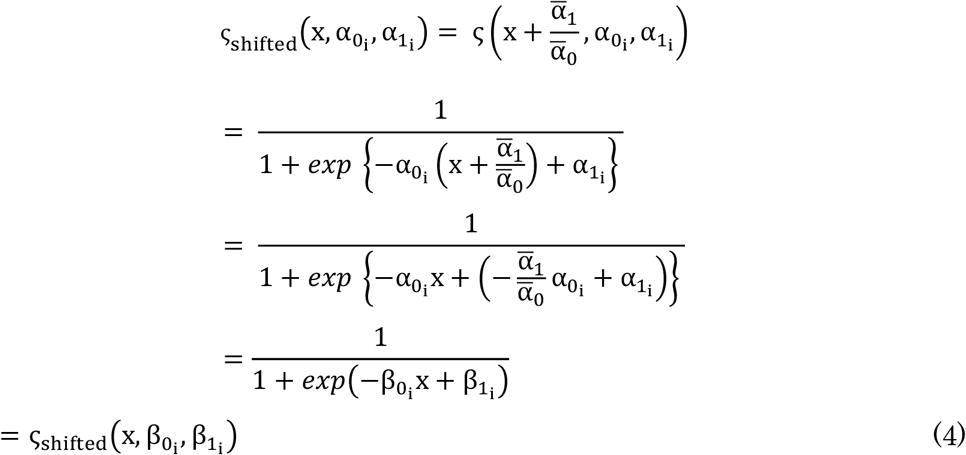

where

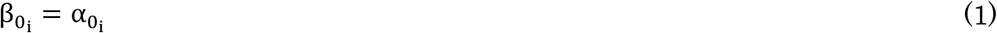

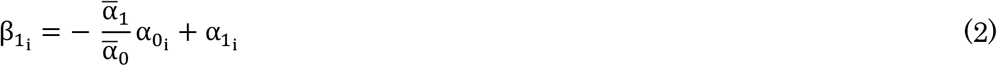

Thus, 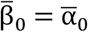 and 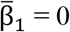 (Fig. 1C).

The mean square error (MSE) between the fitted sigmoid curve and the corresponding sample was calculated. Samples that fell outside the range of the mean ± 3 standard deviations (SDs) in the 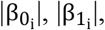, or MSE value for the i^th^ intestine sample were excluded from further analyses, leaving 503 intestine samples.

### Forward passes of neural networks

The three-layer perceptron (control) and INN model had the same structure except for their activation functions. The common structure is written as follows:

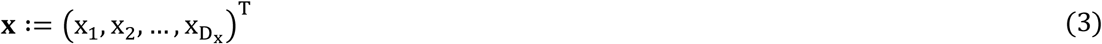

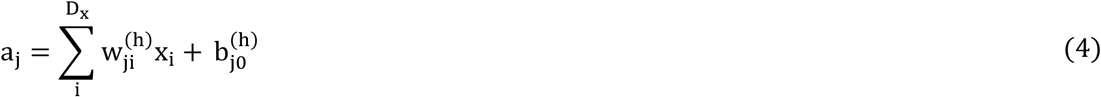

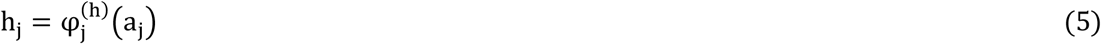

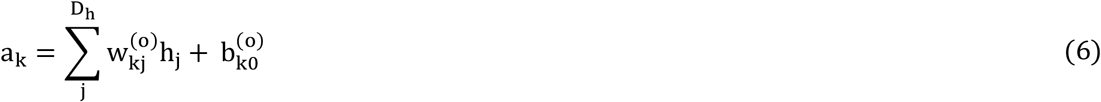

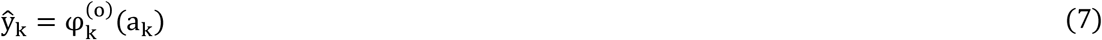

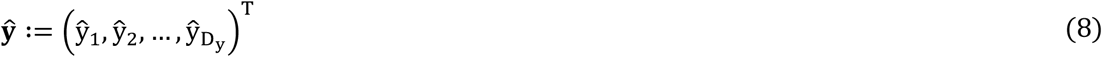

where

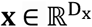: input vector (D_x_: dimension of the input vector)

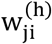: the weight between the i^th^ unit in the input layer and the j^th^ unit in the hidden layer (j = 1, 2, …, 100)

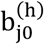: 1, the bias of the j^th^ unit in the hidden layer

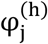: the activation function of the j^th^ unit in the hidden layer

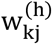: the weight between the j^th^ unit in the hidden layer and the k^th^ unit in the output layer (k = 1, 2, …, D_y_; D_y_ is dimension of the output vector)

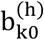: 1, the bias of the k^th^ unit in the hidden layer

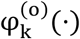: the activation function of the k^th^ unit in the output layer

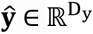: output vector.

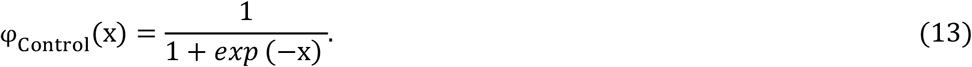

On the other hand, the activation functions of the INN model, 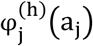 and 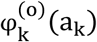, included two random variables B_0_ and B_l_ as parameters:

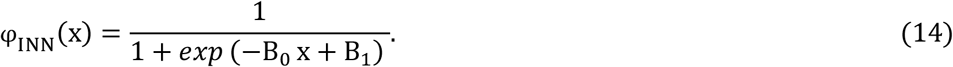

Note that Eq. 13 and Eq. 14 are identical when (B_0_, B_l_) = (1, 0). For each INN activation function, namely, 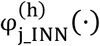 (·) (j = 1, 2, …, 100) and 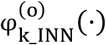 (·) (k = 1, 2, …, D), the random variables (B_0,j_, B_l,j_) and (B_0,k_, B_l,k_) were sampled from 503 sets of experimentally obtained parameters {(β_0_l_, β_l_l_), (β_0_2_, β_l_2_), …, (β_0_503_, β_l_503_)} (Eqs. 1–6).

### Cost function

The cross-entropy loss was adopted as the cost function, which was calculated as follows:

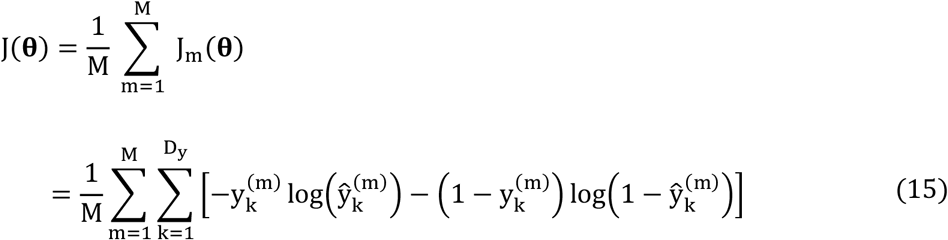

where **θ** represents all the learnable parameters, M denotes the mini-batch size (the amount of training data used in each training loop), J_m_(**θ**) represents the cost function derived from the m^th^ sample in the mini-batch, and y_k_ denotes the k^th^ element of y, which is a one-hot encoded vector of the ground-truth label:

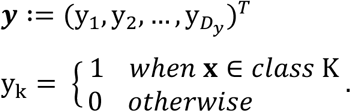

### Backpropagation

The backpropagation method (Rumelhart et al., 1986) was adopted to efficiently calculate the gradient of the cost function, ∇ J_m_(**θ**).

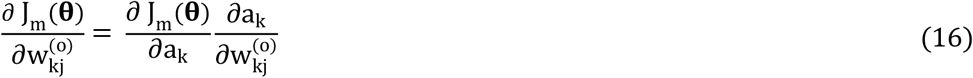

The following notation was introduced:

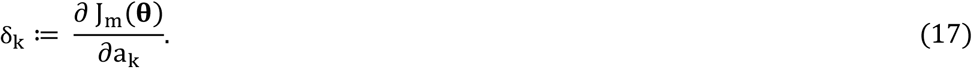

which is analytically defined as:

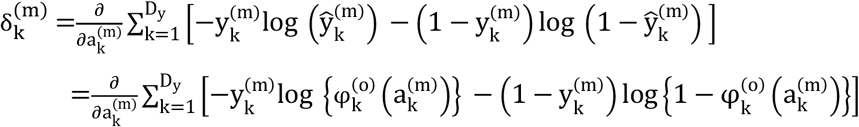

where:

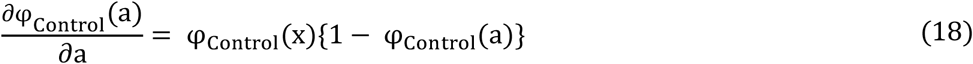

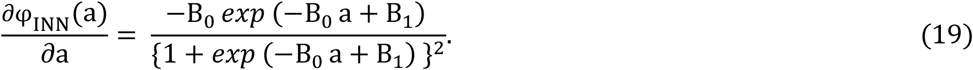

According to Eq. 10, we have

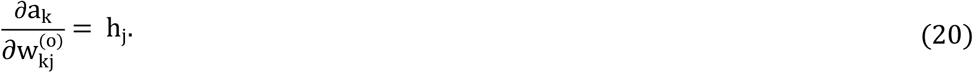

Substituting Eqs. 17 and 20 into Eq. 16, we obtain

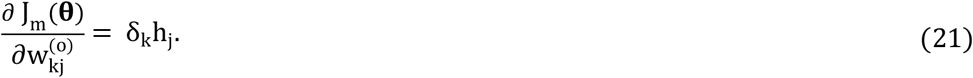

For the δs of the hidden units, the chain rule for partial derivatives is applied to obtain:

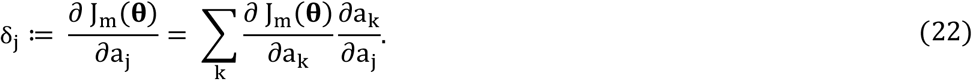

By substituting Eq. 17 into Eq. 22 and using Eqs. 9 and 10, the following backpropagation formula is obtained:

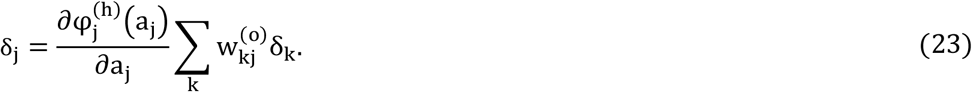

Thus, the gradient of the cost function ∇ J_m_(**θ**) can be calculated using the backpropagation method.

### Weight update

The model was trained using the stochastic gradient descent (SGD) algorithm.

If the error function consists of a sum over data points, J(**θ**) = ∑_n_ J_n_(**θ**), after the presentation of pattern n, the SGD algorithm updates the parameter vector **θ** using the following formula:

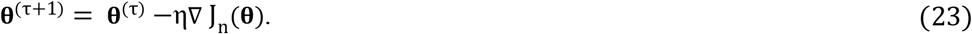

Here, τ denotes the iteration number, and η is the learning rate parameter.

### Classification of handwritten digits

The MNIST database of handwritten digits (LeCun et al., 1998) was used for the experiments in this study. The control and INN models were trained based on 60,000 training data points and evaluated based on 10,000 test data points. Each image was 784 pixels (= 28 columns×28 rows), and the images were fed into models after being flattened into a vector. The mini-batch size was set to 100, and the learning rate was set to 0.05.

### Simulations of INNs

The INN models were simulated by sampling the fitting parameters β_0_ and β_l_. Dummy fitting parameters β_dummy_ = (β_0_dummy_, β_l_dummy_)^T^ were sampled from a bimodal Gaussian distribution, as described below.

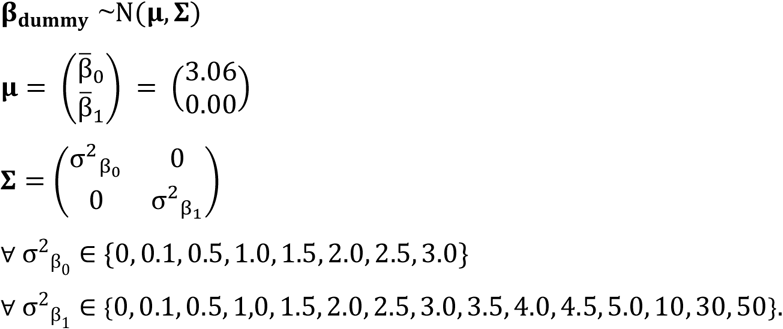

When the slope at the inflection point 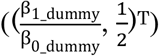 of the simulated sigmoid function (slope: 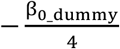 was negative (β_0_dummy_ > 0), **β**_**dummy**_ was resampled. A total of 140 (= 8 × 15) experiments were performed.

### Classification of edible/inedible images

A total of 1,200 images of edible objects and 1,200 images of inedible objects were randomly sampled from https://www.sozaijiten.com. Then, 1,000 images from each class (edible/inedible) were used as training data, and the remaining 200 images from each class were used as test data.

The edible images included images of cooked meals and raw foods such as fruits, vegetables, and fish. The inedible images included images of tools, parts, coins, and other objects.

The images were converted into feature vectors using a pretrained convolutional neural network model (AlexNet; Krizhevsky et al., 2012) based on the Caffe deep learning framework (Jia et al., 2014). Specifically, for every input image, the output of the activation of the fc6 layer (4,096-dimension vector) was extracted.

These feature vectors were fed into the INN model instead of the original images to perform the 2-class classification task. The numbers of units in the input layer, the hidden layer, and the output layer were 4,097 (including a bias unit), 101 (including a bias unit), and 2 (corresponding to two classes: edible or inedible), respectively. The learning rate was set to 0.6.

## Results

### Ileal sample contraction during acetylcholine bath application

After different concentrations of Ach were sequentially applied, the lengths of the isolated ileum tissues decreased. We used an experimental device to measure the isotonic contraction of the intestinal tract (Fig. 1A) and recorded the dose‒response curves of the ilea during bath application of ACh at concentrations ranging from 10^−9^ to 10^−3^ M (Fig. 1B, *n* = 512 tissues from 14 guinea pigs). Each dose‒response curve was obtained from a single tissue and was normalized and fitted to a sigmoid function (Fig. 1C) using two fitting parameters, β_0_ and β_l_ (Eqs. 1–6). The MSE between the experimentally recorded data and the fitted sigmoid curve was calculated for all 512 samples. Data samples for which the |β_0_|, |β_l_| or MSE values exceeded the mean ± 3SD were omitted from the following analysis, and the remaining 503 data points were used to construct the INN model. The distributions of β_0_ and β_l_ are plotted in Fig. 1D and 1E. For β_0_, the sample mean 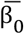 was 3.06, and the sample variance 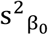 was 1.38; for *β*_l_, the sample mean 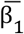 was 0.00 (due to the adjustment), and the sample variance 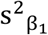 was 3.23).

### Classification of handwritten digits by the INN model

A three-layer perceptron (Control) and an INN with the same structure were built (Fig. 2A), and both models were used to perform a 10-class classification task based on images of the handwritten numbers 0–9 from the MNIST database of handwritten digits (LeCun et al., 1998). During 6,000 rounds of training, the cost of the Control model was lower than that of the INN model (Fig. 2B; at Loop #6,000, Control: 0.558 ± 0.097; INN: 0.923 ± 0.188; *p* < 0.001; *n*s = 40; *U* = 1,556; Mann–Whitney U test). Both models converged, but the sum of the absolute weight update of the Control model was lower than that of the INN model (Fig. 2C; at Loop #6,000, Control: 1.83 ± 0.86; INN: 2.58 ± 0.94; *p* < 0.001; *n* = 40; *U* = 1,182; Mann–Whitney U test). The accuracy of the Control model was higher than that of the INN model (Fig. 2D; at Loop #6,000, Control: 92.3 ± 0.4; INN: 85.7 ± 0.6; Chance: 10%; *p* < 0.001; *n*s = 40; *U* = 0; Mann–Whitney U test).

**Figure 2.**
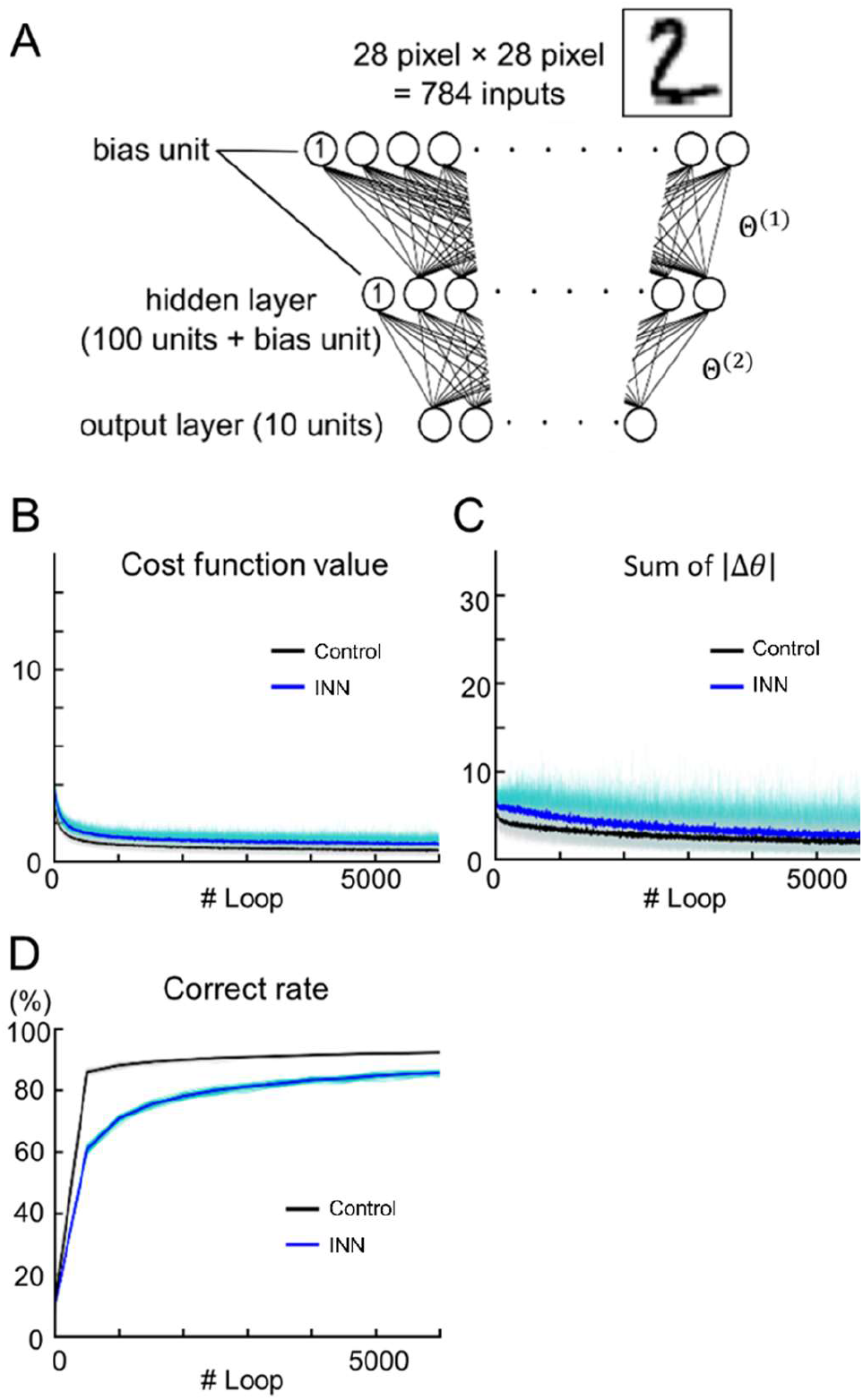
Handwritten number identification using the INN model. (A) A schematic diagram of the three-layer INN model. A conventional perceptron with the same structure was used as the control. (B–D) Output values of test data during training. B, cost function; C, sum of the absolute weight updates; D, classification accuracy. The *blue* and *black* lines represent the INN and control model results, respectively. Each line indicates a single data point (*n* = 32 tuning trials), whereas the thick line indicates the mean.

We also observed the sample variance in the fitting parameters β_0_ and β_l_ (Fig. 1D and 1E). These fluctuations might originate from (i) biological noise and (ii) differences in the experimental conditions. We hypothesized that the variations in β_0_ and 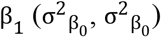 affect the classification accuracy of the model based on the MNIST dataset. To test this hypothesis, we simulated INNs using various values of 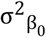 and 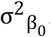. Fig. 3 shows the accuracies of the virtually generated INNs in a pseudocolor map. The highest mean accuracy was 92.3% ± 0.3% for 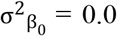 and 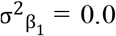, and the lowest mean accuracy was 57.3% ± 5.5% for 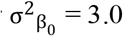 and 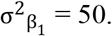.

**Figure 3.**
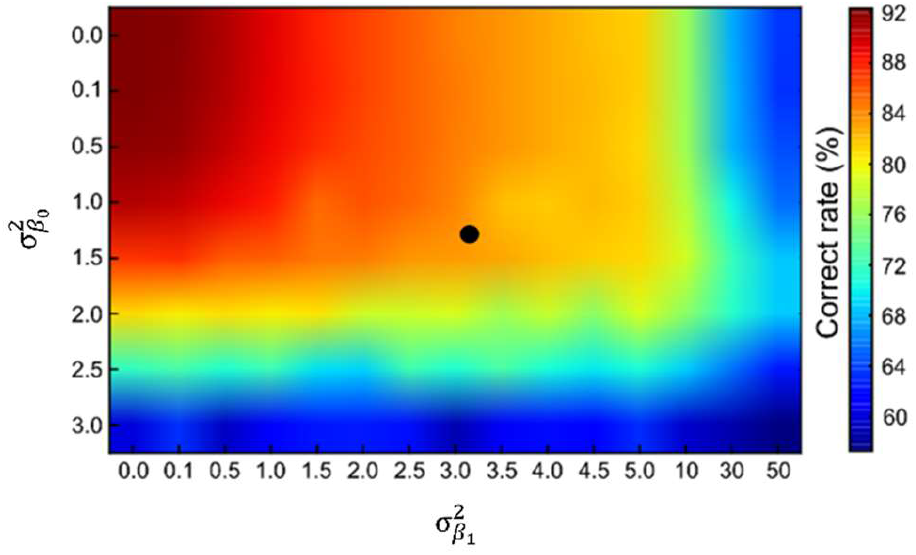
Relationship between INN performance in number identification tasks and parameter variation. Accuracies of INNs as a function of the variance in the hyperparameters β_0_ and 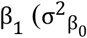 and 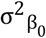). The experimentally obtained sample variance 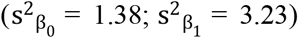 is indicated by the *black* dot.

### Edible/inedible image classification by the INN model

A total of 1,200 images of edible and inedible objects were randomly chosen from the dataset and split into training/test datasets at a ratio of 5:1 (Fig. 4A). The accuracies of the model based on the test data were evaluated during training. At loop #6,000, the accuracy was 88.5% ± 0.9% (Fig. 4; *n* = 32 cross-validations).

**Figure 4.**
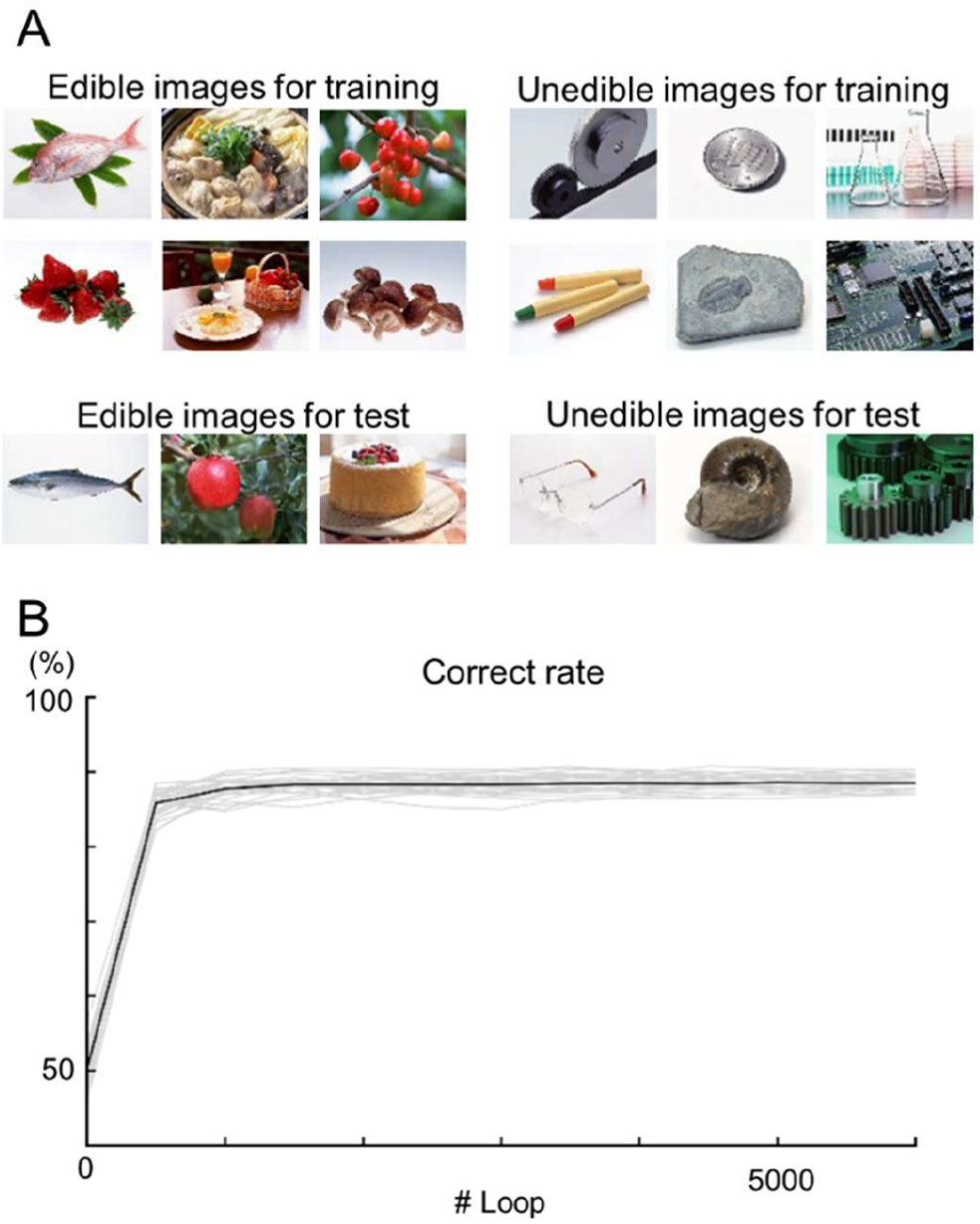
Image classification using the INN model. (A) Examples of edible (*left*) and inedible (*right*) images in the dataset. The top images were used for training, and the bottom images were used for testing. (B) The learning curves of the INN prediction results for the test data. Each *gray* line indicates a single data point (*n* = 32 cross-validation), whereas the *black* line indicates the mean.

## Discussion

In this study, we confirmed that different intestine samples exhibited distinct contraction responses to ACh (Fig. 1B and 1C). Using the response curves, we built an INN model and demonstrated that the INN model could be used to precisely classify handwritten digits. Furthermore, we artificially generated various INN models and found that the INN model with the average activation function, or the three-layer perceptron, achieved the highest accuracy (92.3% ± 0.3%). Finally, the INN model could be used to classify edible/inedible images, achieving a classification accuracy of 88.5% ± 0.9%.

This study presents a novel approach for building neural networks using biological data and demonstrates the potential of the enteric nervous system for developing activation functions for machine learning models. Existing activation functions can be broadly classified into two groups: (i) the sigmoid group, including the logistic sigmoid function and the hyperbolic tangent function, and (ii) the V-shaped group, including the ReLU (Hasnloser et al., 2000; Nair & Hinton, 2010), Leaky ReLU (Maas et al., 2013), PReLU (He et al., 2015), Swish (Ramachandran et al., 2018), and Mish (Misra, 2019) activation functions. However, the use of sigmoid functions has recently decreased because sigmoid functions lead to slow weight updates and vanishing gradients (Bengio et al., 1994) in the backpropagation framework (Rumelhart et al., 1986). Despite this disadvantage, sigmoid functions remain important for biologically inspired network models (Shinozaki, 2021; Zhan et al., 2023), as backpropagation is unlikely to occur in a real brain (Bartunov et al., 2018). Sigmoid curves are known to emerge as cumulative functions for bell-shaped curves, such as normal distributions. The sigmoid curves observed in our pharmacological experiments are regarded as macroscopic expressions of decision-making processes at the biomolecular level in the enteric nervous system because the ACh signaling pathways via muscarinic receptors trigger contractions in smooth muscles (Eglen et al., 1996).

One limitation of our study is the granularity of the model. Our intestine samples were modeled as nodes in an INN, rather than as the network itself. Thus, the developed INN can be viewed as an ensemble model based on intestine samples. This modeling approach was necessitated by the difficulty in recording a large number of individual neurons and receptors. Our model may not necessarily be inaccurate because the central and enteric nervous system exhibit plasticity (*e*.*g*., updated functional connectivity at synapses; Citri & Malenka, 2008) and can be viewed as an ensemble model of different network states at various timepoints. On the other hand, we observed different ACh responsiveness among individual intestine samples (approximately 10^2^ scale). This variation may be caused by biological noise, as indicated by the observed parameters β_0_ and β_l_, which were located in local maxima. Due to these variations, the INN model may have shown different sensitivity to various input data (Fig. 2B) and plasticity (Fig. 2C), which are desirable characteristics in real-world settings, in which the data are often unpredictable and unfixed and balancing the explore-exploit tradeoff is important. To verify this hypothesis, novel datasets and evaluation methods that closely mimic real-world scenarios should be developed to make fair comparisons between *in silico* and biological models.

In conclusion, we successfully developed an INN model using intestine data and demonstrated its potential in performing image classification tasks. This work has important implications in the field of biologically inspired machine learning.

## Acknowledgments

We thank Dr. Hideyoshi Igata for his technical support. This work was funded by JST ERATO (JPMJER1801). Y.I., H.B., and N.H. conceptualized the study; Y.W. performed the experiments and data analysis; Y.W. and Y.I. wrote the original draft; and all authors reviewed and edited the final manuscript.

